# MicroRNAs are differentially altered by ethanol and caffeine consumption in the rat prostate

**DOI:** 10.1101/2025.11.26.690806

**Authors:** M. Martinez, F. S. N. Lizarte, L. F. Tirapelli, D. P. C. Tirapelli, F. E. Martinez

**Author notes:** **Mailing address:** Francisco Eduardo Martinez, PhD, Department of Structural and Functional Biology, Institute of Biosciences of Botucatu (IBB), UNESP - Univ Estadual Paulista, P.O. Box 510, Postal Code: 18618-970, Rubião Júnior, s/n, Botucatu, SP - Brazil.

## Abstract

Simultaneous ingestion of ethanol and caffeine has become popular among young people and teenagers. Intake of caffeine simultaneously with alcohol consumption may increase the risk of alcohol abuse and dependence. Adenosinergic pathways are protective, and caffeine has a non-selective antagonistic role on adenosinergic receptors. Thus, it is assumed that caffeine consumption can interfere with cellular processes inherent to inflammatory mechanisms and may attenuate the onset of inflammation. MicroRNAs play a significant role in cellular and physiological functions and can be altered by consuming ethanol and other substances associated with the modulation of the inflammatory response. This study aimed to identify changes in the expression of microRNAs involved in inflammatory processes and adenosinergic pathways in the prostate, resulting from the interaction between ethanol and caffeine. Male UChB and Wistar rats, aged 5 months, were divided into the following groups (n=15/group): 1. UChB (UChB rats receiving a 10% ethanol solution and water ad libitum); 2. UChB caffeine (UChB rats receiving a 10% ethanol solution with 3g/L caffeine and water ad libitum); 3. Control (Wistar rats receiving only water ad libitum). Animals’ prostate was collected and processed for RT-PCR techniques for miRNAs 155-5p, 146a-5p, 126-3p, - 132-3p, 339-5p. Both ethanol and caffeine consumption modulated the expression of miRNAs related to the inflammatory process and adenosinergic pathways in the tissue of UChB animals. Caffeine may act as a protector of the prostate tissue.

## INTRODUCTION

Alcoholism is a multifactorial syndrome characterized by physical, behavioral, and socioeconomic impairments [1], with its diagnosis, prognosis, and treatment still not fully understood. Alcohol consumption has significantly influenced various societies for thousands of years, playing roles in medicinal, social, and industrial settings, among others. Despite efforts to reduce its prevalence, alcohol remains deeply ingrained in many cultures [2]. Several factors contribute to an individual’s predisposition to ethanol addiction. The risk of developing dependence is influenced by genetic makeup, environmental factors, and neuroadaptations resulting from both acute and repeated drug exposure. Approximately 50% of the risk of alcohol-related disorders is heritable, while the remaining risks are linked to environmental influences. Ethanol alters gene expression and impacts synaptic function, contributing to behaviors such as tolerance, heightened awareness and compulsive consumption, key features of addiction [3].

Caffeine, a xanthine alkaloid, is the most widely consumed psychoactive substance globally. It is found in various foods and beverages, including soft drinks, chocolate, coffee, and tea. Its effects on human behavior have been studied for decades, with ongoing scientific debates regarding its benefits and potential harm [4]. The combined consumption of alcohol and caffeine is increasingly common, particularly among young individuals. Given that ethanol is a central nervous system (CNS) depressant and caffeine a stimulant, many believe that caffeine may counteract or reduce the cognitive and motor deficits caused by alcohol intoxication.

The relationship between coffee consumption and prostate cancer risk has been investigated since the 1960s, yielding inconsistent results. No clear evidence suggests that coffee consumption increases prostate cancer risk [5]. Some studies even associate coffee intake with a reduced risk of prostate cancer, though a causal relationship remains uncertain. Conversely, the combination of atorvastatin and caffeine has been found to regulate apoptosis, migration, and invasion of tumor cells, suggesting a potential strategy for inhibiting prostate cancer growth that warrants clinical evaluation [6]. However, studies on the combined effects of ethanol and caffeine on the prostate are scarce. One study [7] reported no significant association between cigarette use, alcohol intake, coffee, tea, caffeine, and prostate cancer risk.

MicroRNAs (miRNAs) are small, non-coding RNA molecules that regulate gene expression and play essential roles in cellular and physiological processes. Comprising approximately 21 to 23 nucleotides, miRNAs modulate the translation of specific RNAs by binding to complementary regulatory sequences, leading to mRNA degradation or destabilization. Alcoholism is a complex disorder with significant genetic implications. Although the relationship between miRNAs and alcoholism is not fully understood, miRNAs are believed to mediate the effects of ethanol consumption. Identifying serum molecular markers involved in various pathologies, including chronic alcoholism, could improve diagnosis and aid in the development of therapeutic targets. Research suggests that ethanol induces epigenetic modifications, including changes in histone acetylation, methylation, and DNA hypo-or hypermethylation. These findings have led to new research avenues, offering insights into ethanol’s effects on nucleosomes, gene expression, and associated pathophysiological consequences. Ethanol-sensitive miRNAs function as regulatory master switches, potentially influencing the development of tolerance, a hallmark of alcohol addiction. Additionally, other substances of abuse target specific ethanol-sensitive miRNAs, indicating common biochemical mechanisms underlying addiction. Drugs of abuse can modulate gene expression and cellular networks through miRNA regulation. However, the interplay between ethanol and caffeine in this context remains largely unexplored. miRNAs may serve as molecular hubs for drug tolerance and dependence mechanisms. While changes in up to 3% of miRNAs have been observed in ethanol-induced liver disease and teratogenesis models, little is known about miRNA regulation in response to ethanol and its interaction with caffeine [8].

Since the 1940s, researchers have studied rodents with varying preferences for ethanol consumption. Differences in ethanol preference allowed the selection of rats and mice predisposed to low or high ethanol intake, leading to the development of specific animal models. The UChA and UChB rat lines were established in the 1950s at the University of Chile (hence the UCh designation) as models for experimental alcoholism. UChA rats exhibit low voluntary ethanol consumption (0.1–2 g ethanol/kg body weight/day), whereas UChB rats have high voluntary intake (5–7 g ethanol/kg body weight/day) [9]. The UChB strain, a genetically selected pure line of Wistar rats, serves as a valuable model for studying the genetic, biochemical, physiological, nutritional, and pharmacological effects of ethanol. These rats voluntarily consume ethanol, closely mimicking chronic human alcohol dependence.

This study aimed to investigate alterations in microRNA expression associated with inflammatory processes and adenosinergic pathways in the prostate, resulting from ethanol and caffeine consumption in UChB rats.

## MATERIAL AND METHODS

### Animals and Selection of Ethanol Consumption

Thirty adult rats of the UChB variety, male, five months old, from the Vivarium of the Department of Structural and Functional Biology, Institute of Biosciences, UNESP - Campus of Botucatu-SP, from an important International Cooperation with the University of Chile, and 15 rats were used. Wistar, male, five months old, from the Animal Facility of UFSCar/São Carlos-SP. The animals were housed individually in polypropylene cages with bedding and kept under controlled conditions of luminosity, humidity, and temperature (12-hour light/dark cycles, 55 ± 5% humidity, and 20 - 25 °C). They had ad libitum access to commercial chow (Purina) and water. Three experimental groups were formed (N=15): (1) a group of UChB rats with voluntary consumption of 10% ethanol (water + ethanol = 7 g of ethanol/kg body weight/day), (2) a group of UChB rats with voluntary consumption of 10% ethanol (water + ethanol = 7 g of ethanol/kg body weight/day) plus caffeine (300 ml/l), and (3) a control group of Wistar rats with voluntary consumption of water ad libitum, used as a comparison for the UCh rats that naturally consume alcohol and for those exposed to both alcohol and caffeine. The day of birth for the litter was designated as day zero. Weaning occurred at 21 days of age, with the pups housed in groups of two to four animals to prevent stress from social isolation. At 50 days of age, the rats were housed individually and received regular handling appropriate for the species. At 65 days of age, the animals were given two bottles: one with water ad libitum and the other with a 10% ethanol solution, which were periodically alternated. After 15 days of assessing the ingestion of the 10% ethanol solution, the animals that consumed an average of approximately 7 g of ethanol/kg body weight/day during this period were selected and standardized to the UChB lineage. From the end of the selection period until euthanasia (80 to 150 days of age), ethanol consumption was monitored every 7 days. Three groups of 10 rats each were then established: UChB, UChB + caffeine (300 mg/l), and Control (Wistar). The UChB + caffeine group continued voluntary consumption of the 10% ethanol solution plus caffeine for 55 consecutive days (from 95 to 150 days of age). The experimental protocol adhered to ethical principles in animal research, as outlined by the Brazilian School of Animal Experimentation (SBCAL/COBEA). All rat experiments were conducted in compliance with the National Institutes of Health guide for the care and use of laboratory animals. The protocol also followed the ethical standards set by the National Council for the Control of Animal Experimentation (Brazil) and the Committee on Ethics in the Use of Animals (CEUA) at the Federal University of São Carlos (UFSCar).

### Macroscopy

The ventral lobes of the prostates were dissected, with or without the assistance of a stereoscopic microscope, collected, and then immediately fixed in a liquid nitrogen solution.

### Material Processing for Molecular Biology

Fragments of the ventral lobes of the prostates from five animals per group were collected. The ventral lobes of the prostates were stored in liquid nitrogen and kept in a - 80 ºC freezer. Next, according to the protocols below, these materials were prepared for analysis of MicroRNAs - PCR in real-time.

### MicroRNAs 155-5p, 146a-5p, 126-3p, 132-3p, 339-5p in the ventral prostatic lobe

Five animals per group were anesthetized using ketamine chloride (50 mg/mL) (Ketalar, Parke-Davis) and xylazine (20 mg/mL) (Rompum, Bayer) in doses of 90 and 10 mg/kg, respectively, which were administered simultaneously via the peritoneal cavity. After anesthesia, the ventral lobe of the prostate was collected and frozen in liquid nitrogen. RNA extraction was performed using Trizol™ reagent (Invitrogen, USA). For complementary DNA (cDNA) synthesis, reverse transcription was carried out using the High-Capacity cDNA Reverse Transcription kit (Applied Biosystems) according to the manufacturer’s instructions. The microRNAs 155-5p, 146a-5p, 126-3p, 132-3p, and 339-5p were analyzed using the TaqMan Assay-on-Demand system, which includes oligonucleotides and probes obtained from Applied Biosystems. Amplification was performed in a final volume of 10 µL, consisting of 5 µL of TaqMan Master Mix (Applied Biosystems), 0.5 µL of each specific probe, and 4.5 µL of cDNA. All reactions were carried out in duplicate and analyzed using real-time PCR detection equipment (7500 Real-Time PCR System, Applied Biosystems). Data were continuously collected during PCR and analyzed with ABI-7500 SDS software. Statistical analysis of gene expression was conducted using the Shapiro-Wilk normality test and One-way ANOVA. GraphPad Prism version 4.0 for Windows (GraphPad Software) was used for analysis, with significance set at P < 0.05.

## RESULTS

### MicroRNAs 126-3p, 132-3p, 146a-5p, 155-5p and 339-5p in the ventral prostatic lobe

#### Ethanol has influenced the expressions of prostate miRNAs

Ethanol ingestion did not influence statistically the expressions of 126-3p, 132-3p, 146a-5p, 155-5p, and 339-5p miRNAs in the ventral prostatic lobe. Animals that ingested ethanol had similar miRNA levels to animals in the control group. However, a non-statistically significant increase was observed in miRNAs 132-3p, 155-5p, and 339-5p after ethanol consumption **(Fig. 1)**.

**Fig. 1.**
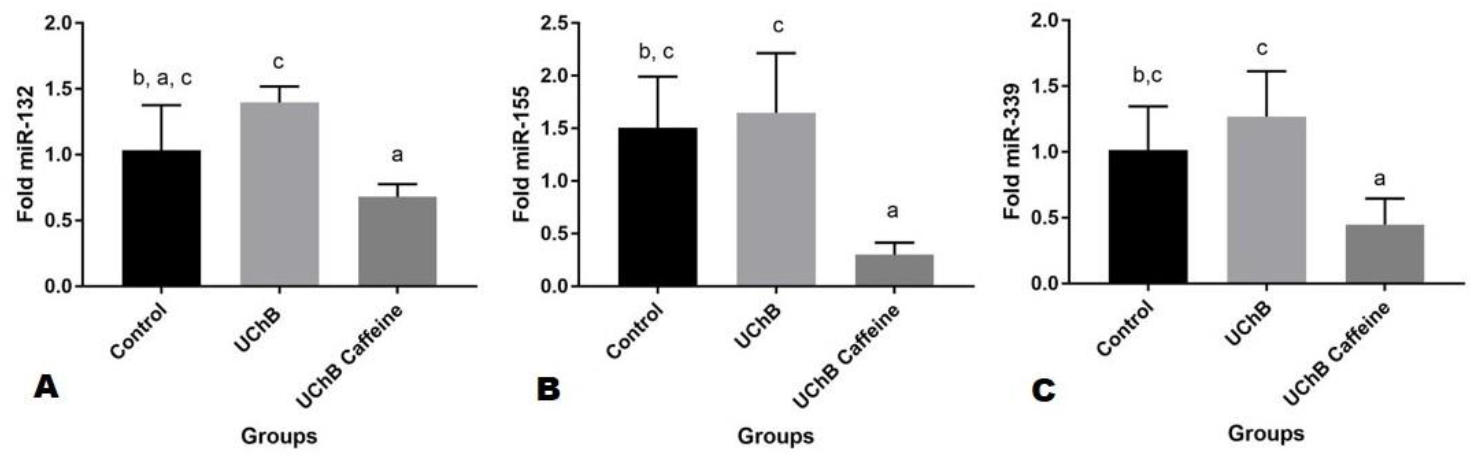
Shows the mean (± standard deviation) expression of MicroRNAs 132-3p **(A)**, 155-5p **(B)**, and 339-5p **(C)** in the prostate of rats. Data are presented as mean ± standard deviation. Five animal samples were considered per group. Different letters indicate statistically significant differences between groups. *p < 0.05, **p < 0.01, ***p < 0.001.

#### Caffeine decreased the level of prostate miRNAs

Caffeine consumption influenced the levels of miR-126-3p, miR-132-3p, 146a-5p, 155-5p, and 339-5p in the UChB caffeine group compared to the Control group. Caffeine was able to reduce miRNAs 126-3p, 132-3p, 146a-5p, 155-5p, and 339-5p to lower levels than the control group. **(Figs. 1, 2)**.

**Fig. 2.**
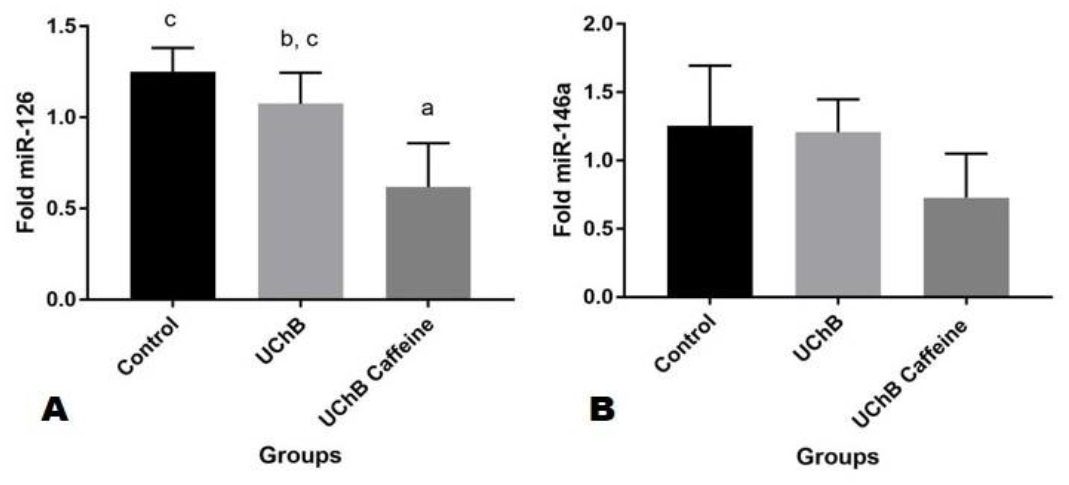
Shows the mean (± standard deviation) expression of MicroRNAs 126-3p **(A)** and 146a-5p **(B)** in the prostate. Data are presented as mean ± standard deviation. Five animal samples were considered per group. Different letters indicate statistically significant differences between groups. *p < 0.05, **p < 0.01, ***p < 0.001.

## DISCUSSION

The consumption of ethanol and caffeine significantly altered miRNA levels in both prostate tissues. Few studies have examined caffeine’s modulation of serum miRNAs, with most research focusing on its effects on tissues [10]. Caffeine is a potent antioxidant, proposed to inhibit oxidative stress-induced changes in miRNA transcription. The abnormal elevation of miRNAs can silence multiple genes, impairing protein translation. Caffeine counteracts this effect by suppressing the elevation of toxic miRNAs and preventing target gene silencing. Studies suggest that miRNAs can either mitigate or mediate drug-related behavioral effects, highlighting their role in neurobehavioral development following fetal exposure to substance abuse [11, 12]. Drugs of abuse can alter specific miRNA levels, downregulating target genes and leading to abnormal cellular function. Ethanol-induced changes in up to 3% of miRNAs have been documented in models of ethanol-induced liver disease and teratogenesis [13].

The increase in miRNA-155-5p enhances the secretion of pro-inflammatory cytokines and nitric oxide [14]. miR-155-5p is a well-known pro-inflammatory miRNA essential for macrophage and T-cell responses. In macrophages, miRNA-155 is stimulated by toll-like receptor (TLR) ligands and cytokines such as interferon-γ (IFN-γ) [15]. A study [16] showed that miRNA-155 deficiency protects against ethanol-induced increases in pro-inflammatory cytokines and NF-κB production. High miR-155 expression interferes with the cell cycle, promoting cancer cell proliferation, migration, invasion, and immune evasion, particularly in liver cancer. The present study supports these findings, as ethanol consumption slightly increased miR-155-5p expression, while caffeine ingestion attenuated this increase to levels comparable to the Control group.

MiRNAs, including miR-146a-5p, contribute to resolving inflammation. Overexpression of miR-146a-5p in microglial cells increases IL-10, a key regulator of inflammation [17]. In astrocytes treated with IL-1β, higher miR-146a expression negatively regulates inflammation by reducing IL-6 and COX-2 levels. miR-146a-5p directly targets genes in MyD88 signaling, including IRAK-1 and TRAF-6, critical for NF-κB activation via TLRs [18]. In this study, ethanol consumption in UChB rats did not increase miR-146a expression. However, caffeine consumption reduced miR-146a expression below Control levels, suggesting that the tissue does not require additional inflammatory suppression. The simultaneous increase in miR-155-5p and miR-146a-5p is commonly observed in inflammation, indicating an active tissue response [15].

Chronic ethanol consumption increased miR-132 expression in this study. A previous study [19] demonstrated that elevated miR-132 levels upregulate glutamatergic receptors GluA1, GluN2A, and GluN2B, suggesting ethanol modulates not only inflammatory but also structural proteins. miR-132 is a key regulator of hepatic lipid homeostasis, functioning in a context-dependent manner through the suppression of multiple targets with cumulative synergistic effects. Ethanol + caffeine consumption reversed this expression to Control levels.

A significant decrease in miR-126-3p expression was observed in the UChB caffeine group. MiR-126-3p plays a key role in angiogenesis, vascular homeostasis, and the innate immune response [20]. During inflammation, miR-126 suppresses reactive oxygen species production in endothelial cells by modulating HMGB1 expression [21]. These findings highlight that miRNA expression in specific tissues does not always correlate with systemic effects.

MiR-339-5p expression was altered in the UChB and UChB caffeine groups. Previous studies indicate that miR-339-5p functions as a tumor suppressor, inhibiting prostate cancer cell proliferation, migration, and invasion [22]. Another study [23] found that RNA circ_0086722 competitively binds to miR-339-5p, alleviating its suppressive effects on STAT5A, and thereby promoting prostate cancer progression. These findings provide insight into prostate tumor progression and potential therapeutic targets.

Caffeine regulates miRNA expression and, as a xanthine alkaloid, possesses strong antioxidant properties. It inhibits oxidative stress-induced transcriptional changes in DNA [24]. Tissue response is mediated by adenosinergic receptors, with adenosine acting as a key regulator through G protein-coupled receptors. Many immune cells express these receptors and respond to adenosine’s modulatory effects [25]. However, little is known about how caffeine attenuates cell loss. It is hypothesized that caffeine delays degenerative processes primarily by reducing inflammation [26], as demonstrated in this study through its regulation of miRNAs.

Caffeine also influences miRNA expression through post-transcriptional mechanisms. Alternative splicing (AS) of pre-mRNA selectively joins exons to produce diverse mRNA variants from a single gene [27]. AS can alter untranslated regions (UTRs) or introduce premature codons, resulting in mRNA variants subject to regulation affecting translation efficiency, localization, or stability [28]. One regulatory mechanism, nonsense-mediated decay (NMD), ensures accurate gene expression by degrading non-productive mRNAs, including those from AS [29]. Caffeine inhibits NMD, promotes alternative 3’ UTR splicing, and downregulates target miRNAs, thereby modulating post-transcriptional processes [24].

Both ethanol and caffeine exert antagonistic effects on the central nervous system. Caffeine intake should be carefully controlled, as no universal safe daily limit exists. The consumption of up to 400 mg per day (approximately four cups of coffee) by adults (70 kg) and non-pregnant women is considered non-harmful, whereas pregnant women should limit intake to less than 300 mg daily [4].

MicroRNAs have emerged as valuable bioinformatics tools and non-invasive diagnostic and therapeutic targets. They not only predict disease occurrence but also enable genetic-level treatments. Advancements in drug delivery systems are gradually achieving precision therapy by regulating intracellular miRNA levels [30].

The increasing consumption of caffeine and ethanol underscores the importance of understanding their interactions with the adenosine receptor system. Further research is needed to clarify the effects of chronic simultaneous ethanol and caffeine ingestion, providing new avenues for genetic studies. The findings of this study open perspectives for future genetic analyses to elucidate differences in susceptibility between UChB, UChA, and other Control groups regarding ethanol and caffeine consumption.

## STUDY LIMITATION

While our results provide novel insights into the molecular interactions between ethanol and caffeine in the prostate, limitations must be acknowledged. The study focused solely on microRNA expression without assessing downstream protein targets or functional outcomes.

## CONCLUSIONS

The interaction between caffeine and ethanol affected miRNA expression in complex ways, and more broadly cellular biology, potentially having implications for health, the treatment of substance abuse-related diseases, and future genetic research in this field.

1. **Alteration of miRNAs in Response to Ethanol and Caffeine Consumption**: The simultaneous consumption of ethanol and caffeine significantly altered the levels of several miRNAs in prostate tissues, suggesting that these substances play an important role in modulating the expression of miRNAs that regulate numerous functional processes.
2. **Effects of Ethanol and Caffeine on Inflammatory miRNAs**: miR-155-5p and miR-146a-5p are involved in inflammatory processes. The increased expression of miR-155-5p is associated with the secretion of pro-inflammatory cytokines, while miR-146a-5p is known to help resolve inflammation. However, caffeine intake helped mitigate the inflammatory effects caused by ethanol, suggesting a protective effect of caffeine against inflammation.
3. **Regulation of miR-132 and miR-126-3p**: Chronic ethanol consumption increased the expression of miR-132, which may be related to the modulation of glutamatergic receptors and structural proteins. On the other hand, caffeine was able to reverse this expression. miR-126-3p, which plays an essential role in angiogenesis and the maintenance of vascular epithelial integrity, was reduced in the UChB + caffeine group, suggesting that caffeine may modulate the vascular and immune response.
4. **Caffeine and Its Potential Regulation of miRNAs**: Caffeine has a strong antioxidant potential and may interfere with the transcription of miRNAs induced by oxidative stress. Additionally, caffeine can regulate alternative splicing and the degradation of non-productive mRNA, impacting various post-transcriptional processes that influence the stability and translation of genes. This effect may contribute to attenuating cellular damage caused by ethanol.
5. **Interaction Between Ethanol and Caffeine**: Although caffeine and ethanol have opposing effects on the central nervous system, their interactions need to be better understood. The simultaneous consumption of these substances may have complex effects, influencing gene expression and cellular processes in contradictory ways, which could affect the risk of dependence or other issues related to the abuse of these substances.
6. **Perspectives for New Genetic Studies**: The study suggests that genetic research could provide valuable insights into how different animal strains, such as UChB and UChA, respond to ethanol and caffeine consumption. This could help identify genetic factors associated with susceptibility to substance abuse and provide new directions for therapy and prevention.
7. **Therapeutic Potential of miRNAs**: MiRNAs are presented as an emerging tool for precision diagnostics and therapies. They can be used to predict disease development and as targets for therapeutic interventions, aiming to regulate miRNAs to treat diseases related to substance consumption.

## AUTHOR CONTRIBUTION STATEMENT

M M, and F E M conceived the study, performed experiments, analyzed, and finished data, and wrote the paper. F S N L, and M M performed experiments. L F T, D P C T, L G A C, and F E M provided training to perform the experiments and intellectual input for the experimental design and data analysis. All authors contributed to editing the paper.

